# Evidence that glycopolymer transferases promote peptidoglycan hydrolysis in *Bacillus subtilis*

**DOI:** 10.1101/2025.02.26.640348

**Authors:** Jennifer D. Cohen

## Abstract

Most bacteria are encased in a rigid cell wall peptidoglycan (PG) meshwork. Cell growth requires the activities of both PG synthases and PG hydrolases that cleave bonds within the meshwork enabling its expansion. PG hydrolase activity must be carefully regulated to prevent excessive damage to this protective layer leading to catastrophic lysis. Here, I provide evidence for a novel type of regulation mediated by lipid-linked glycopolymer precursors. The Gram-positive bacterium *Bacillus subtilis* encodes two functionally redundant PG hydrolases, LytE and CwlO, that are required for growth. Here, I demonstrate that loss of LytR-CpsA-Psr (LCP) enzymes, which enzymatically transfer lipid-linked glycopolymers onto PG, leads to a requirement for *lytE* for growth. Genetic analysis suggests that this requirement is mediated by the accumulation of these membrane-anchored precursors, where they may interfere with PG hydrolase activity. These results are consistent with models in which polymer transfer influences the position or timing of PG hydrolysis.

## Introduction

Bacteria must continuously synthesize and remodel their cell envelopes to enable growth. The primary load-bearing layer of the cell envelope is the cell wall peptidoglycan (PG), a single macromolecule of glycan strands crosslinked by short peptides (Warth and Strominger, 1971). Too little cleavage of PG restricts expansion of growing bacterial cells; however, too much hydrolysis leaves cells vulnerable to lysis (Weidel and Pelzer, 1964). Thus, the hydrolase that degrade PG must be tightly regulated. Altering their regulation to allow too much or too little PG degradation may be an attractive target for novel antibiotics. However, due to a wide range in number and type of PG hydrolases, how PG cleavage is regulated, and whether it is coordinated with PG synthesis, remains poorly understood (Vollmer et al., 2008).

In the widely-studied Gram-positive bacterium *Bacillus subtilis*, two secreted PG-cleaving enzymes, LytE and CwlO, are redundantly required for PG expansion and are heavily regulated (Bisiccia et al., 2007; Margot et al.,1998). In wild-type cells, neither *lytE* nor *cwlO* is required for growth; however, when both are mutated, cells fail to elongate due to a lack of PG cleavage (Bisiccia et al., 2007; Hashimoto et al., 2012). LytE contains three glycan-binding LysM domains that allow it to bind to the PG layer (Steen et al., 2003; Chen et al., 2008). In contrast, CwlO is present in a membrane complex containing the AAA ATPase FtsEX and a pair of small membrane proteins, SweD and SweC. Loss of *ftsEX* or *sweDC* completely abolishes CwlO function, rendering *lytE* essential for growth (Domínguez-Cuevas et al., 2013; Meisner et al., 2013; Brunet et al., 2019). *B. subtilis* monitors CwlO activity on a whole-cell level via the WalRK two-component signaling pathway, which senses low CwlO activity and in response increases expression of PG hydrolases, including CwlO and LytE, and decreases inhibitors of CwlO and LytE activity (Dobihal et al., 2019; Bisiccia et al., 2007; Bisiccia et al., 2010; Salzberg et al., 2013; Dobihal et al., 2022). However, loss of *lytE* and *cwlO* does not inhibit PG synthesis (Meisner et al., 2013). Instead, if there is a connection between LytE and CwlO activity and PG synthesis, it must occur via a different mechanism.

One possible regulator of the LytE and CwlO hydrolases are the glycopolymers that are covalently attached to PG in Gram-positive bacteria. Loss of the most abundant polymer, wall teichoic acid (WTA), yields rounded, slow-growing cells that have high levels of LytE, indicating that these cells likely have a defect in PG structure (D’Elia et al., 2006; Kasahara et al., 2016). *B. subtilis* also synthesizes many other PG-anchored polymers: capsule, minor teichoic acids, teichuronic acids, and exopolysaccharide (EPS), as well as polymers likely to be produced by several so-far-unstudied gene operons within the *B. subtilis* genome (Armstrong et al., 1959; Baddiley 1970; Marvasi et al., 2010; Branda et al., 2006; Kearns et al., 2005; Hamon and Lazazzera 2001; Morikawa et al., 2006; Freymond et al., 2006; Bhavsar et al., 2004; Janczura et al., 1961; Shuster et al., 2019). Polymers are strong candidates to communicate the status of PG synthesis to membrane-bound protein complexes because these lipid-linked polymers are transferred onto newly-synthesized PG, and thus their abundance in the membrane might serve as a proxy for localized PG synthesis (Schaefer et al., 2017). However, whether these lipid-linked glycopolymer precursors influence PG assembly or impact LytE and CwlO activity is not known.

In this work, I provide evidence for an inhibitory relationship between the accumulation of membrane-anchored polymers and CwlO activity in *B. subtilis*. These lipid-linked polymers are transported from the inner leaflet to the outer leaflet of the cytoplasmic membrane and then transferred to PG by membrane-anchored LCP (LytR-CpsA-Psr family) enzymes (Kawai et al., 2011). LCP enzymes are widely conserved among both Gram-positive and Gram-negative bacteria (Hübscher et al., 2008). In Gram-positive bacteria, LCP enzymes transfer a wide variety of undecaprenyl-phosphate (C55-P)-linked polymers onto nascent PG (Schaefer et al., 2017; Chan et al., 2014; Eberhardt et al., 2012; Harrison et al., 2016; Sigle et al., 2016; Baumgart et al., 2016; Wu et al., 2014). In this work, I show that depletion of LCP enzymes that accumulate lipid-linked precursor caused phenotypes consistent with decreased CwlO activity. My data suggest that CwlO activity was partially restored in cells lacking enzymes that are predicted to function in the synthesis of glycopolymers. These results are consistent with a model in which buildup of membrane-anchored glycopolymer precursors impair CwlO activity. These results suggest that PG-linked polymers can act as regulators of PG hydrolysis in *B. subtilis*.

## Results

### The *B. subtilis* LCP enzymes function together to promote growth and maintain cell shape

To investigate the hypothesis that loss of polymers from PG may impact PG assembly and degradation, I generated a set of *B. subtilis* strains lacking the three genes encoding LCP enzymes (called *tagT, tagU,* and *tagV* in *B. subtilis*). *tagT, tagU, and tagV* encode transmembrane proteins (TagTUV) with extracellular LCP domains that transfer polymers from their membrane-embedded undecaprenyl-phosphate (C55-P) carriers onto PG (Schaefer et al., 2017; Li et al., 2020) (Fig 1A). Loss of all three enzymes completely eliminates the abundant polymer WTA from the PG layer *in vivo* (Kawai et al., 2011). I found that strains lacking all 3 LCP paralogs (Δ*tagTUV*) were viable, but marginally so; cells were rounded and grew extraordinarily slowly, similar to reports of mutants lacking WTAs (D’Elia et al., 2006) (Fig S1A). The extremely slow growth of the *ΔtagTUV* mutant was similar to what was previously reported for a LCP mutant in *Staphylococcus aureus* (Over et al., 2011) and only slightly healthier than the phenotype reported previously for the *ΔtagTUV* mutant in *B. subtilis* (Kawai et al., 2011). To create a more tractable strain for determining the roles of *tagTUV* in cell growth, I introduced an IPTG-regulated allele of *tagV* or *tagV-His* at an ectopic locus in the *ΔtagTUV* strain. I separately generated a strain in which the promoter of the operon that includes *tagV* (*yvyE-tagV*) was fused directly to *tagV*. All three *tagV* alleles complemented the slow growth phenotype of the *ΔtagTUV* strain indicating they were functional (Fig 1B, 1C, S1B). The strains harboring the IPTG-regulated *tagV* alleles grew at a slower rate in the absence of IPTG (Fig 1B). Importantly, these cells had a severely rounded shape, similar to the *ΔtagTUV* strain (Fig 1C).

**Figure 1.**
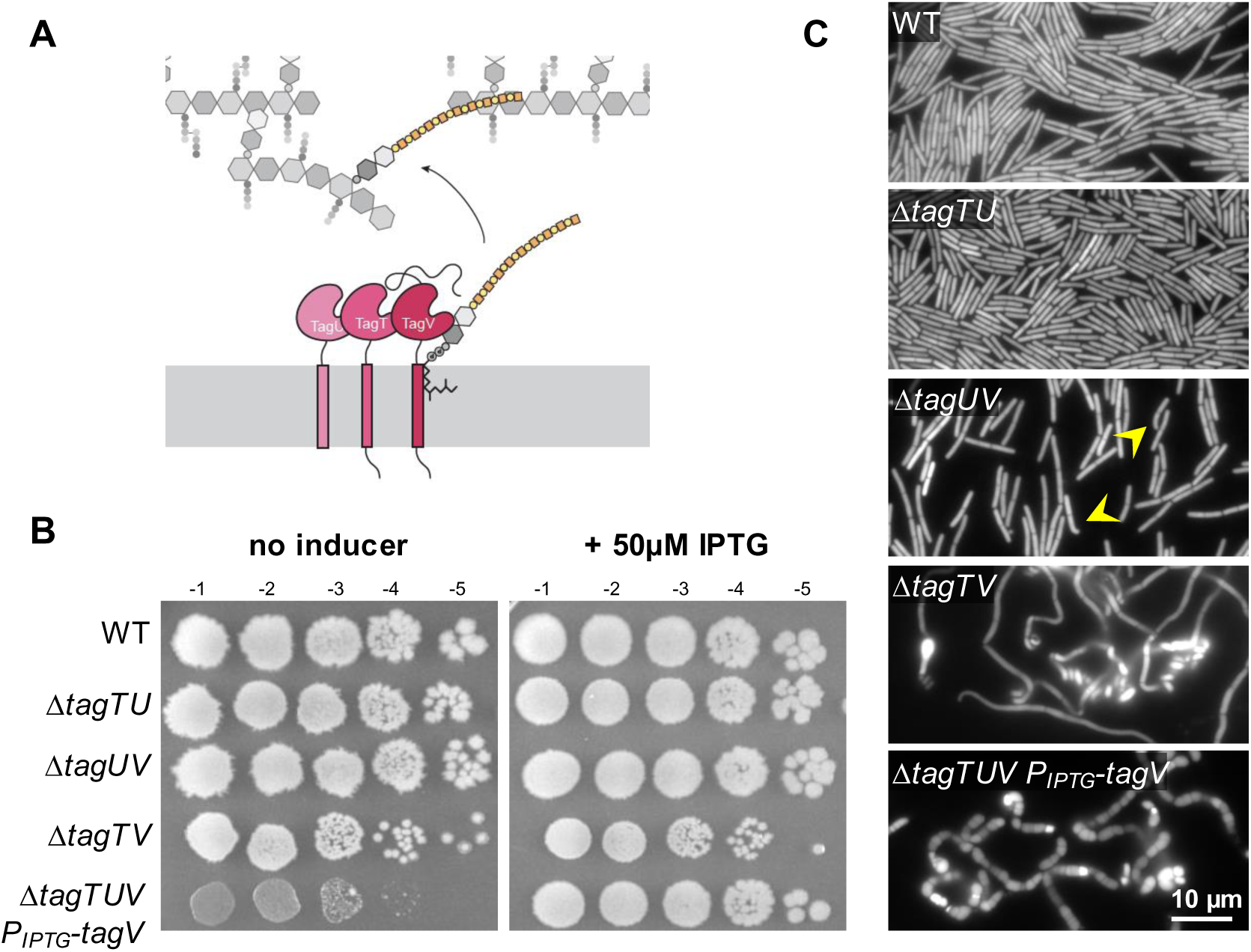
*B. subtilis* LCP enzymes act synthetically. A) Cartoon depicting TagTUV (pink) transferring polymers (yellow) from the periplasmic surface of the membrane onto PG (gray). B) Spot dilutions of the indicated strains in the presence or absence of IPTG. Triple deletion mutants lacking all three B. subtilis LCP genes *(ΔtagTUV*) express *tagV* under an IPTG-inducible promoter and grow very slowly in the absence of inducer. C) Fluorescence images of the indicated strains grown in LB media. WT cells are rod-shaped. Cells lacking *tagV* and *tagU* show slight morphology defects (arrowheads). Cells lacking *tagV* and *tagT* are coiled. Cells lacking all three LCP genes grown without inducer are severely rounded.

Mutants lacking only two of the LCP enzymes revealed that *tagV* is the most important LCP enzyme for growth. While single Δ*tagT, ΔtagU, ΔtagV* or double *ΔtagTU* mutants had a wild-type morphology and grew well on LB agar plates, a subset of the *ΔtagUV* mutant cells had bent shapes indicative of defects in the cell envelope (Fig 1C). Finally, *ΔtagTV* double mutants formed slightly small colonies on agar (Fig 1B) and had abnormal, twisted morphologies (Fig 1C). This analysis indicates that TagT, TagU, and TagV function redundantly, but that TagV and TagT are the more important of the three paralogs for cell growth and morphology. The *in vivo* importance of these enzymes could be due to differences in function or expression level.

### Cells lacking LCP enzymes require LytE for growth

In the course of my characterization of the *ΔtagTUV* mutant, I investigated synthetic interactions with mutations in the PG hydrolases *lytE* and *cwlO*. These experiments revealed that cells lacking *lytE,* but not *cwlO,* were strongly impaired in growth in absence of *tagTUV* and *tagTV* (Fig 2A,B and S2). Since the only functional PG hydrolase in the *ΔlytE* strain is CwlO, these results suggest that CwlO activity depends on the LCPs.

**Figure 2.**
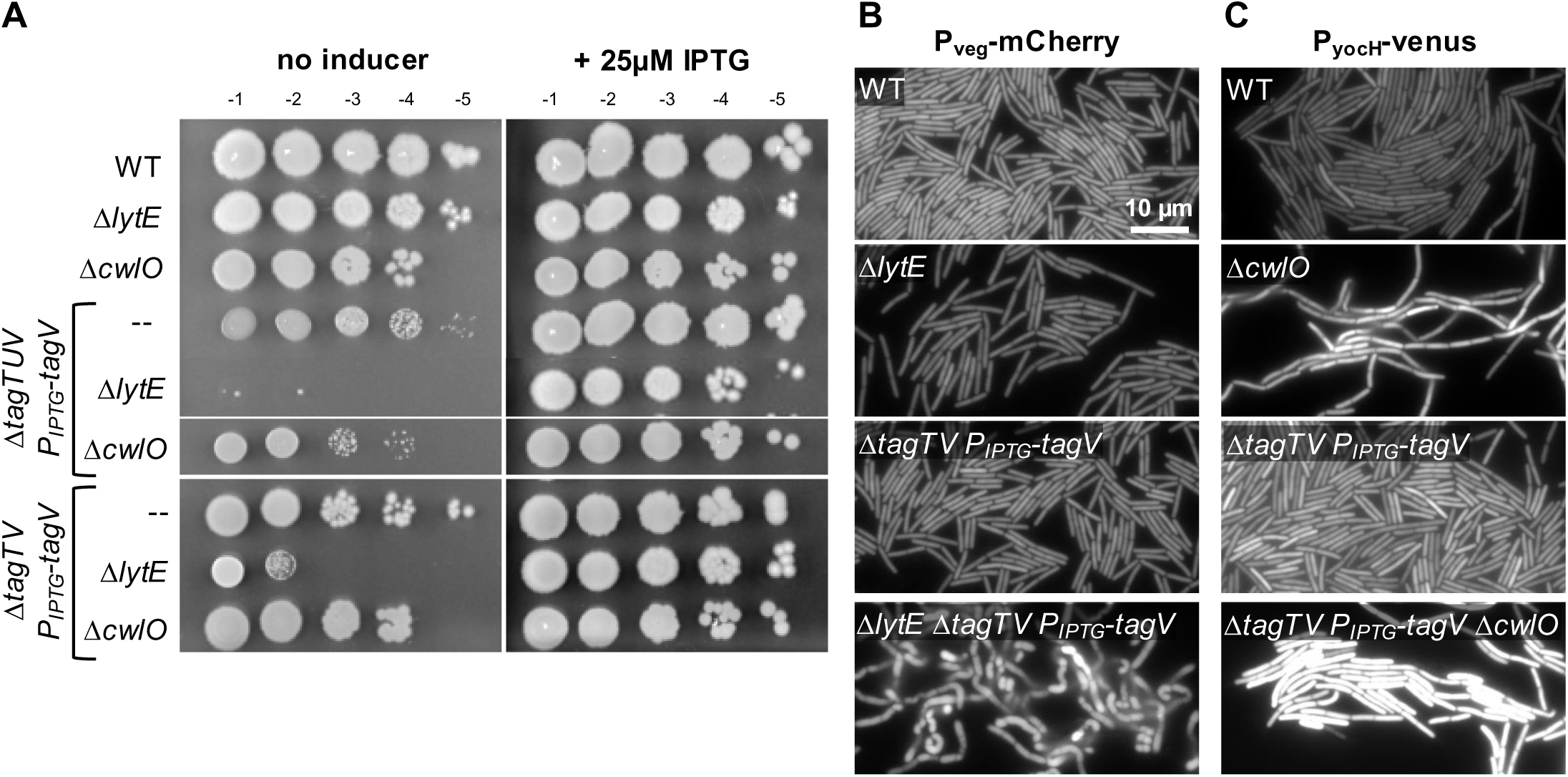
LCP genes promote *cwlO* activity. A) Spot dilutions of the indicated strains in the presence or absence of IPTG. *ΔtagTUV P_IPTG_-tagV* grow very slowly in the absence of inducer, but do not grow at all when *lytE* is deleted. *ΔtagTV P_IPTG_-tagV* cells grow at normal rates on LB-agar without IPTG, but are severely growth limited in the absence of *lytE.* Neither strain shows decreased growth when *cwlO* is deleted. B) Constitutively-expressed, cytoplasmic mCherry reveals that *ΔlytE ΔtagTV* mutants have severely abnormal morphology. *ΔtagTV P_IPTG_-tagV* cells have normal morphology, perhaps due to low levels of leaky expression of *tagV*. C) *ΔcwlO* and *ΔtagTV* mutants have similarly high expression of *PyocH-venus*, indicating higher levels of WalRK signaling. These levels are increased further in double mutants.

To further examine whether CwlO activity is affected by low levels of LCP enzymes, I performed a series of genetic experiments. First, I analyzed the morphology of LCP depletion mutants that are also lacking *lytE*. Because *ΔtagTUV* mutants already have a severely rounded shape, whereas *ΔtagTV* mutants are elongated but curved and filamented (Fig 1C), I compared a *ΔtagTV* double mutant *to a ΔtagTV ΔlytE* triple mutant. The triple mutant had a rounded morphology and growth arrest that closely mirrored the phenotype of *ΔlytE* mutants depleted of CwlO cofactors. These findings are consistent with the idea that CwlO activity is impaired in *ΔtagTV* mutants (Meisner et al., 2013; Brunet et al., 2019). To further investigate whether CwlO activity is decreased in *ΔtagTV* mutants, I investigated if WalRK activity is increased in *ΔtagTV* mutants. WalRK is activated when CwlO activity is reduced, and WalRK activity can be measured using a WalR-responsive transcriptional reporter in which the *yocH* promoter is fused to YFP (Dobihal et al., 2019; Bisiccia et al., 2007; Bisiccia et al., 2010; Salzberg et al., 2013; Dobihal et al., 2022). Using this reporter, I found that *ΔtagTV* mutants have increased WalRK activity that is similar to cells lacking *cwlO*, supporting the model that CwlO activity is reduced in the absence of two of the LCPs (Fig 2C). However, the *ΔtagTV ΔcwlO* triple mutant had even higher WalRK activity than either *ΔtagTV* or *ΔcwlO* mutants, raising the possibility that reduced LCPs not only impairs CwlO activity but also impacts the activity of other cell wall hydrolases that contribute to WalRK activation.

### LCP enzymes do not promote *cwlO* activity through known pathways

Do TagTUV act through the FtsEX-SweDC complex that regulates CwlO activity (Meisner et al., 2013; Brunet et al., 2019)? To test if CwlO is released from this membrane complex in *ΔtagTUV* mutants, I expressed *tagV-His* at various levels in a *ΔtagTUV* mutant (Fig 1, S1). I generated protoplasts lacking the cell envelope, following the protocol of Brunet et al., 2019 and determined the levels of CwlO, FtsEX, and SweDC by immunoblot. The data in Figure 3 show that the levels of all 5 of these proteins were unchanged regardless of *tagV* expression level. These data indicate that TagTUV are not involved in anchoring CwlO or stabilizing the FtsEX complex (Fig 3A). Next, I tested whether a mutation in *ftsE* that bypasses the requirement for *sweDC* in a strain lacking LytE also bypasses the requirement for *tagTUV* (Brunet et al., 2019). This *ftsE* allele did not suppress the growth defect of the Δ*tagTV ΔlytE* triple mutants (Fig 3B). Though the nature of this suppressor allele is unknown, these data suggest that TagTUV do not directly impact FtsEX activity. Finally, AlphaFold3 did not predict an interaction between TagV and FtsEX or SweDC (Fig S3), arguing against a model in which these proteins form a complex *in vivo* (Abramson et al., 2024). Attempts to test the requirement for enzymatic activity of TagV by mutating its active site in a manner similar to TagU in Kawai et al., 2011 failed to generate a stable protein, possibly due to changes in the active site of TagV relative to TagU (Li et al., 2020) (data not shown). More research is necessary to determine if TagTUV interact with the FtsEX-SweDC-CwlO complex *in vivo*. However, the data so far suggest that *tagTUV* is not required for expression or stability of FtsEX-SweDC, that TagTUV do not impact CwlO interaction with the FtsEX-SweDC complex, and cannot be suppressed by a mutation in *ftsE* that suppresses the requirement for *sweDC*.

**Figure 3.**
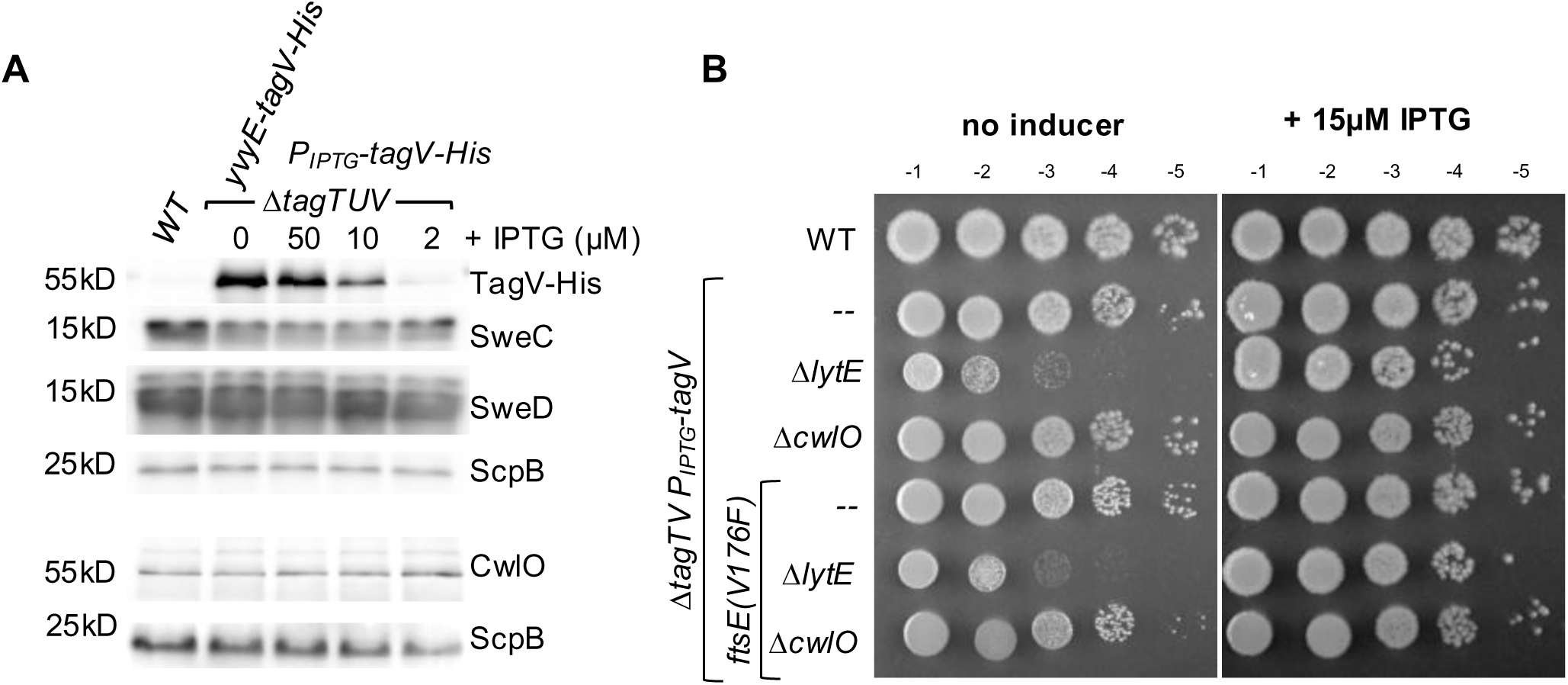
***tagTUV* do not promote *cwlO* activity via the FtsEX-SweDC pathway.** A) Western blot of labeled TagV and endogenous SweDC and CwlO. ScpB, loading control. Levels of SweDC and CwlO were stable as TagV levels were depleted. B) Spot dilutions of the indicated strains in the presence or absence of IPTG. An *ftsE* mutation that suppressed Δ*sweDC; ΔcwlO* growth defect had no effect on *ΔtagTV*; Δ*cwlO*.

### PG-attached polymers and the depletion of C55-P do not impact CwlO activity

How might TagTUV influence CwlO activity? In the absence of *tagTUV,* the lipid-linked polymers are not transferred onto PG. Thus, polymers are lost from the PG; the carrier lipid C55-P is trapped; and lipid-linked polymers accumulate in the periplasmic space (Fig 4). Each of these defects could individually impact CwlO or FtsEX activity, leading to three models for how TagTUV influences CwlO. In the first model, loss of polymers attached to the PG inhibits the ability of CwlO to cut the meshwork. In the second model, trapped C55-P due to accumulation of WTA precursors impacts CwlO activity. In a third model, the accumulation of lipid-linked polymers impairs CwlO activity. These models are not mutually exclusive, nor are they the only possible models. The following experiments were designed to determine which of these models most likely reflects the underlying mechanism.

**Figure 4.**
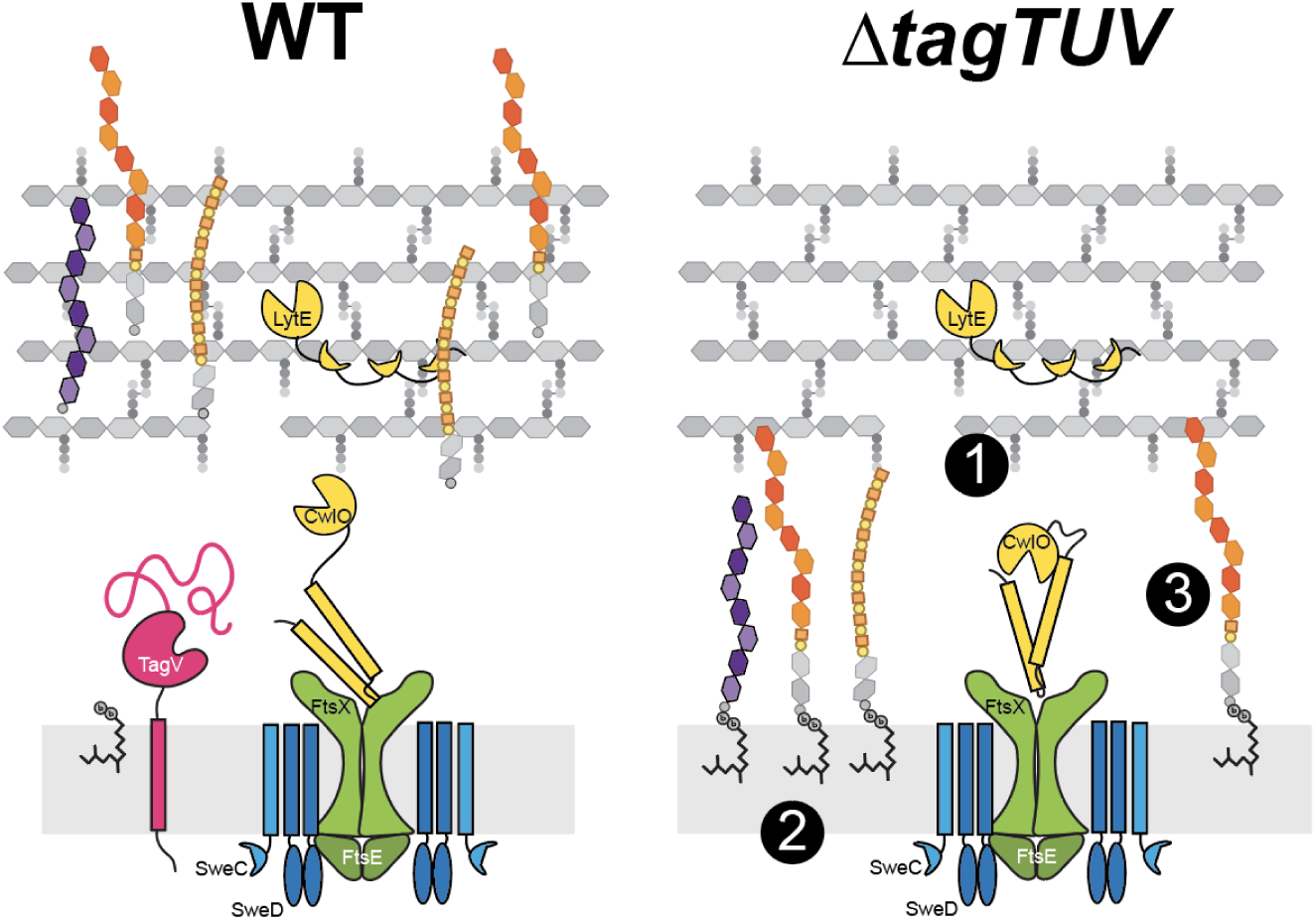
Models for TagTUV promotion of CwlO activity. Model 1: The loss of PG-attached polymers inhibits CwlO activity. Model 2: The loss of the free lipid carrier, undecaprenyl phosphate (C55-P), and/or its accumulation in the membrane inhibits CwlO activity. Model 3: The accumulation of one or more polymers at the membrane inhibits CwlO.

To test the first model, in which polymers attached to PG are required for CwlO, I constructed a series of deletions of key genes or operons that synthesize polymers. LCP enzymes can transfer many types of polymers from the carrier lipid C55-P onto PG. While the best-known substrate transferred by LCP enzymes is the polyglycerol phosphate polymer WTA precursor (Baddiley 1970), *B. subtilis* produces a number of other polymers that are transferred by LCP enzymes, including EPS, teichuronic acid, minor teichoic acids and capsule (Fig S4). A single enzyme, encoded by *tagO,* performs the first committed synthesis step of WTAs, minor teichoic acids, and potentially teichuronic acids (Soldo et al., 2002). Low levels of TagO expression did not yield synthetic lethal phenotypes in *ΔlytE* mutants, strongly suggesting that PG-attached glycopolymers (WTAs, minor teichoic acids, and teichuronic acids) do not impact CwlO activity (Fig 5A).

**Figure 5.**
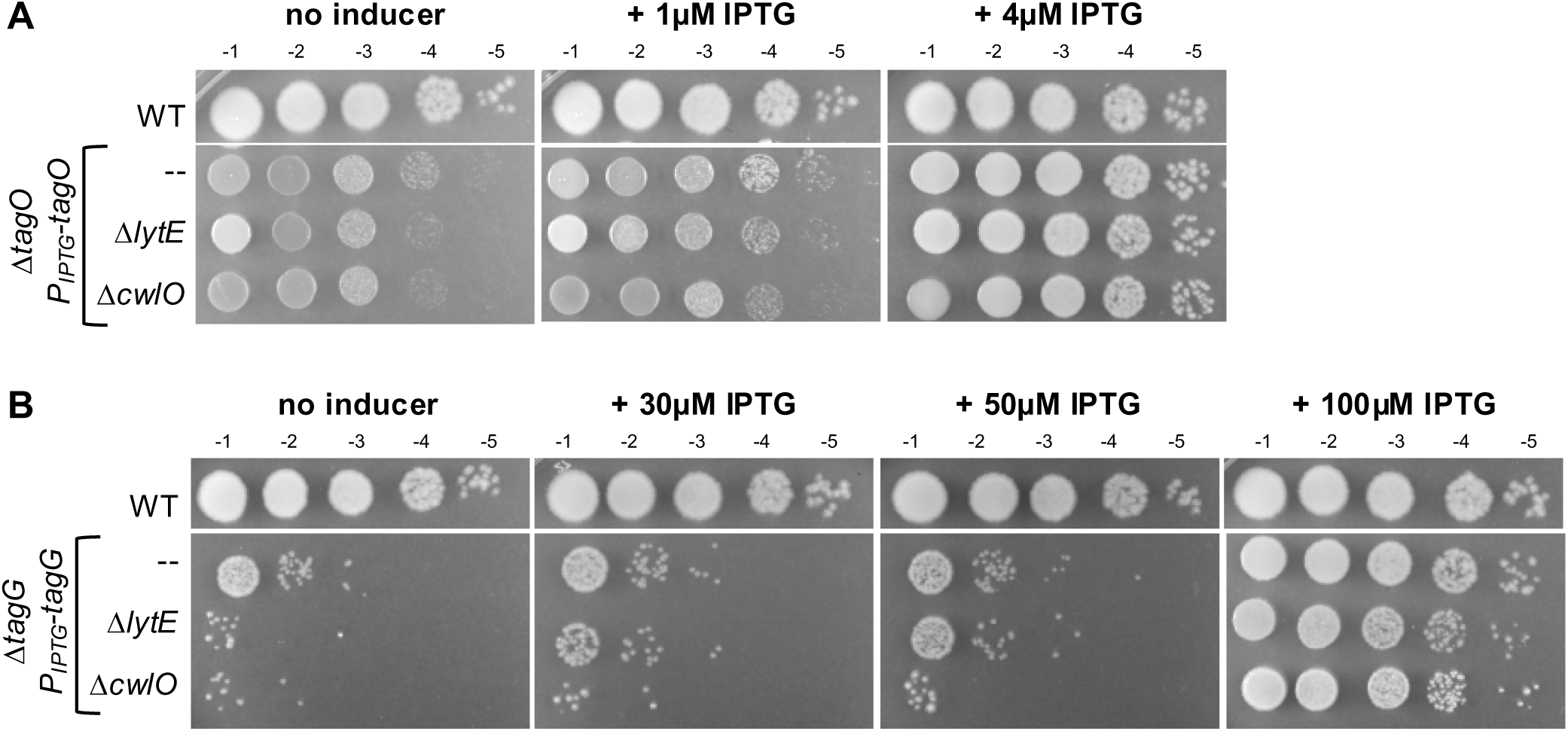
WTAs and trapped C55-P loss do not activate D,L endopeptidases. A) Spot dilutions of the indicated strains with varying concentrations of IPTG. No synthetic effect is observed in *ΔlytE* or *ΔcwlO* mutants depleted for *tagO.* B) Spot dilutions of the indicated strains with varying concentrations of IPTG. Depletion of *ΔtagG* does not cause a synthetic defect with either *ΔlytE* or *ΔcwlO*.

Next, I tested if other PG-attached polymers other than WTAs might influence ClwO activity. Deletions of the operons encoding EPS and capsule caused no growth defect in a *ΔlytE* mutant, suggesting that PG-attached polymers are not critical for CwlO activity (Fig 6A, B). While I cannot rule out the possibility that PG-attached polymers do not promote CwlO activity in combination to cleave PG, my data suggest model 1 is unlikely to explain the loss of CwlO activity in the LCP mutants.

**Figure 6.**
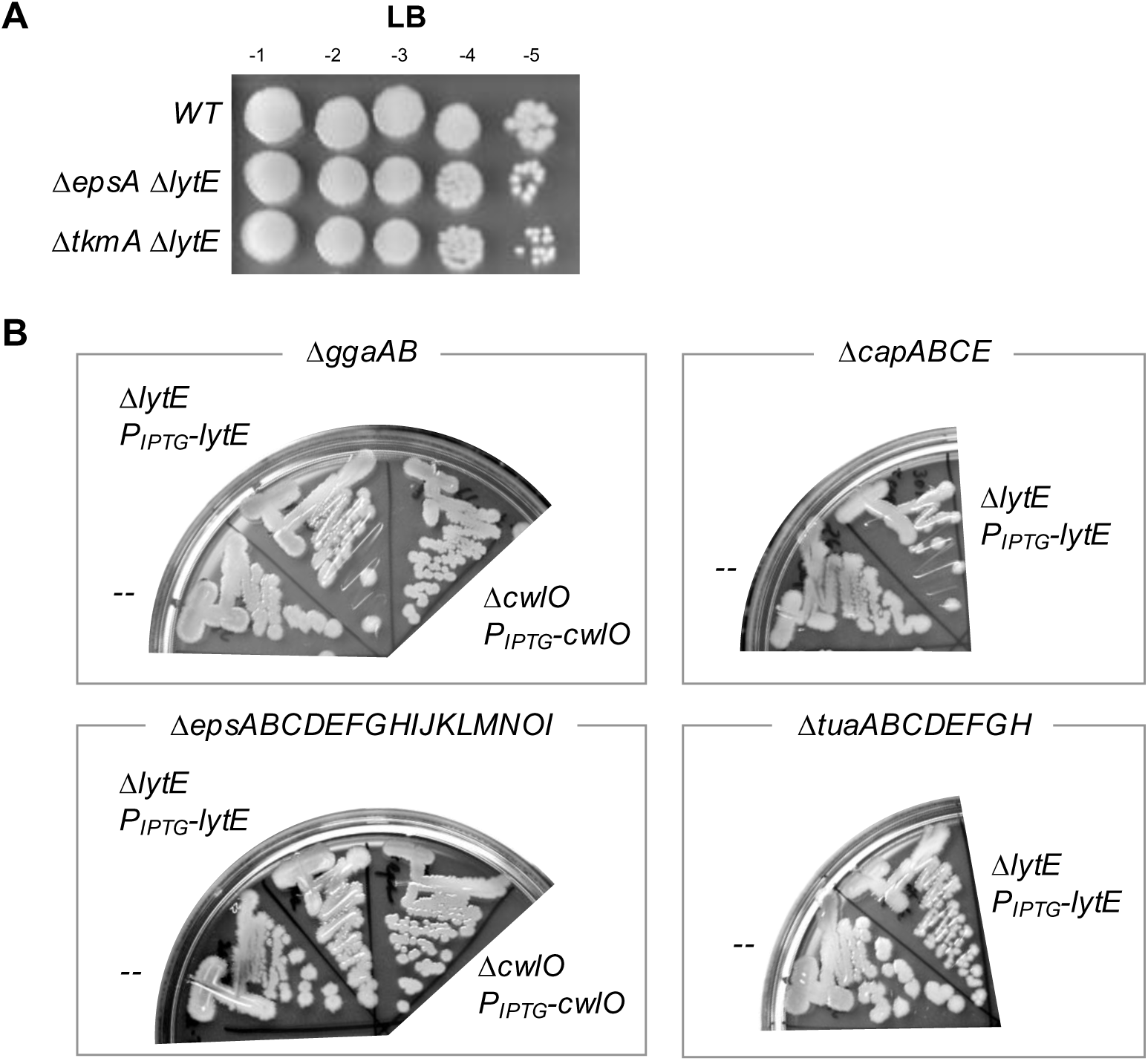
Other polymers do not activate D,L endopeptidases. A) Spot dilution of the indicated strains on LB-agar. No genetic interactions were observed between *lytE* and *epsA* or *tkmA.* B) Streaks of the indicated strains onto LB-agar without IPTG. Genotypes include the loss of the operon encoding polymer synthesizing genes at the top of each box with additional mutations in *lytE* or *cwlO* as indicated.

To test the second model, in which the carrier lipid C55-P on which glycopolymers are built become trapped when these polymers are not transferred, I trapped C55-P-linked WTA precursors in the inner leaflet of the cytoplasmic membrane. To do this, I depleted *tagG*, which is required for transport of the WTA precursor across the cytoplasmic membrane (Fig S4) (Lazarevic and Karamata, 1995; Schirner et al., 2011). Loss of *tagG* leads to a buildup of WTA precursors in the inner leaflet of the cytoplasmic membrane, which is lethal because it traps the essential lipid carrier C55-P (Hancock et al., 1976; Hancock and Baddiley, 1976; Roney and Rudner, 2024). Deletion of *tagG* using an IPTG-inducible allele, did not have synthetic phenotype with Δ*lytE*, indicating that reduction in the carrier lipid is unlikely to influence CwlO activity (Fig 5C).

### The Wzz homolog *tkmA* may inhibit *cwlO* in the absence of LCP enzymes

Finally, I tested if lipid-linked glycopolymer precursors inhibit CwlO activity by mutating key genes required for polymer synthesis in a *ΔtagTV ΔlytE* background. In *B, subtilis,* three homologous genes, *capA, epsA*, and *tkmA* encode Wzz, or PCP, domain proteins that are thought to cleave polymers as they are synthesized to regulate polymer length (Whitfield et al., 2020). *capA* encodes a protein that interacts with capsule-synthesizing enzymes, while *epsA* and *tkmA* function in exopolysaccharide synthesis in biofilm-producing strains of *B. subtilis* (O’Reilly et al., 2023; Kiley and Stanley-Wall, 2010; Gerwig et al., 2014; Gao et al., 2015) (Fig S4). *tkmA* also influences the expression of genes that produce additional polymers by activating the transcription factor PtkA (Shuster et al., 2019; Jers et al., 2010). The broad roles in regulating polymers and their non-essentiality made these genes attractive as tests for whether altered synthesis of polymers could alter CwlO activity in *tagTUV* mutants. Loss of one of these genes, *tkmA,* slightly suppressed the growth of *ΔtagTUV* mutants (Fig 7A). *ΔtkmA* partially suppressed the growth defect of *ΔtagTV ΔlytE* (Fig 7A). Lastly, Δ*tkmA* also suppressed the high levels of WalRK signaling as assessed by *yocH* promoter expression (Fig 7B). Based on these data, I propose that buildup of TkmA-synthesized glycopolymers and C55-P-linked WTA precursors inhibit CwlO activity. When the level of these precursors drops due to their transfer to PG, CwlO activity increases.

**Figure 7.**
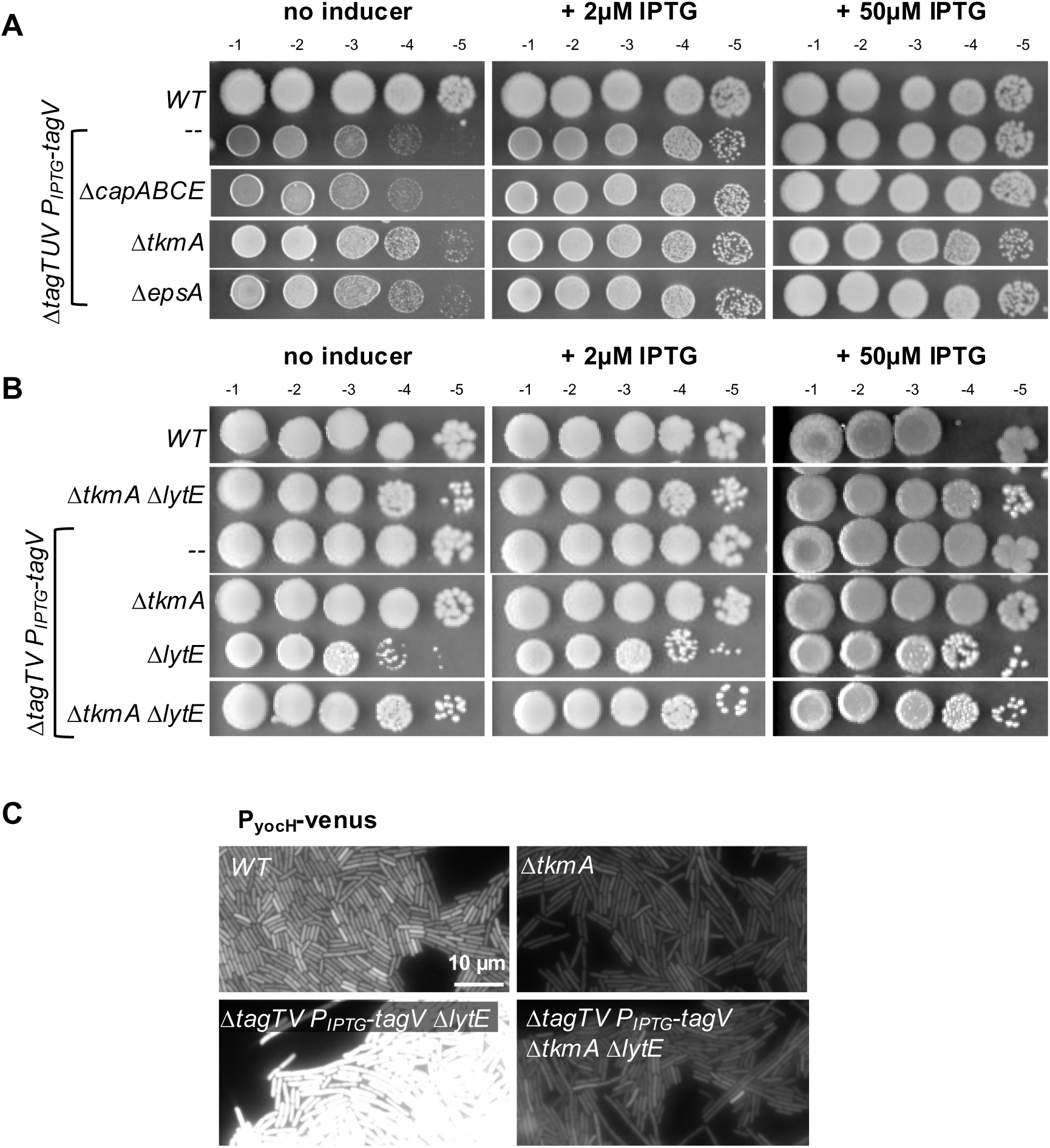
***tkmA* inhibits *cwlO* in the absence of *tagTV*.** A) Spot dilutions of the indicated strains with varying levels of IPTG. Loss of *tkmA* slightly suppressed the growth defect of *ΔtagTUV* mutants expressing low levels of *tagV*. B) Spot dilutions of the indicated strains with varying levels of IPTG. Loss of *tkmA* restored growth of *ΔtagTV ΔlytE* mutants (bottom row). C) Fluorescence images of the indicated strains grown in LB media. *ΔtkmA* also suppressed high WalRK activity as assessed by *yocH* promoter activity.

## Discussion

Altogether, this work raises the possibility that the transfer of lipid-linked glycopolymers from the membrane onto PG may help coordinate PG synthesis and hydrolysis. In the absence of the LCP enzymes TagTUV, cells grow slowly and become rounded, similar to cells lacking TagO, which is required for the synthesis of many of the C55-P linked precursors. My genetic analysis strongly suggests that in *ΔtagTUV* mutants, CwlO activity decreases and LytE activity becomes essential. The *ΔtagTV ΔlytE* synthetic growth defects is largely suppressed when the transmembrane protein, TkmA, is absent. The function of TkmA is currently ill-defined. I propose it contributes to glycopolymer synthesis both as part of an enzymatic protein complex, and indirectly via a cytoplasmic interaction with the kinase and transcription factor PtkA. I hypothesize that in its absence fewer polymers are synthesized, alleviating inhibition of CwlO activity in the *ΔtagTV* mutant.

A major and largely unanswered question in the field of Gram-positive bacterial growth is how these organisms expand their thick PG matrix without triggering excessive PG hydrolysis. Indeed, PG hydrolases were originally identified as autolysins, responsible for cell lysis during stress (Vollmer et al., 2008). The experiments reported here are consistent with a model in which, under unperturbed growth, lipid-linked polymers are transferred to nascent PG by LCP enzymes and removal of these polymers from the membrane relieves inhibition of PG hydrolases. These hydrolases can then cleave PG in regions where nascent synthesis is most active, enabling cell wall expansion. Furthermore, the locally transferred polymers could plug holes in the PG matrix that would otherwise endanger cell integrity. If PG synthesis stops, nascent PG will not be available, and LCP enzymes will not transfer polymers from the membrane. Those membrane-bound polymers may act as a signal that PG synthesis is stalled to locally inhibit PG-cleaving enzymes. Once nascent PG synthesis resumes, LCP enzymes could initiate transfer of polymers, allowing PG hydrolysis to resume. Thus, the work reported here suggests that polymer transfer might function to coordinate PG synthesis with hydrolysis. Future experiments should determine whether polymer type or number at the membrane changes in *ΔtkmA* and *ΔtagTV* mutants. These experiments should also test whether WalRK signaling and/or PG synthesis machinery is suppressed by the presence of a polymer at the membrane surface. Lastly, as it remains possible that other polymers are involved in besides those whose synthesis is promoted by *tkmA,* future experiments should test if C55-linked WTA similarly inhibits CwlO activity via depletion of TagO.

Consistent with the work described in this paper and the corresponding model that polymer transfer to PG coordinates PG synthesis and hydrolysis, published work on other Gram-positive bacteria show that PG-anchored polymers are key mediators between PG synthesis and degradation. For example, in *Streptococcus pneumoniae*, the key PG hydrolase LytA binds to glycopolymers at the cell surface. When these glycopolymers are transferred from the cell membrane onto the PG layer, LytA is also transferred to the PG layer, where it can digest PG (Flores-Kim et al., 2019). Similarly, in *Clostridioides difficile, Staphylococcus aureus,* and *B. subtilis*, WTAs prevent binding of specific enzymes that degrade PG, concentrating these enzymes at specific regions such as the septum where they separate newly formed cells (Wu et al., 2016; Schlag et al., 2010; Frankel and Schneewind, 2012; Yamamoto et al., 2008). The *S. aureus* hydrolase Atl may be sequestered at the cell surface by WTAs that have not been transferred (Tiwari et al., 2018). Similarly, in *Lactococcus lactis,* high galactose media causes changes in the composition of membrane-anchored polymers, leading to increased binding of the PG hydrolase AcmA to those polymers and less PG hydrolysis (Steen et al., 2008). Although in each species the mechanism is different, a common theme is that glycopolymers regulate PG hydrolysis.

Although *cwlO* and *lytE* are coordinately essential for PG hydrolysis, they appear to have starkly different regulation. LytE is thought to be released from the cell surface to bind PG glycans via its three LysM domains, where it may cleave outer layers of PG (Steen et al., 2003; Chen et al., 2008). In contrast, CwlO remains tethered to the cell membrane by FtsEX, where it would seemingly have access to only the most membrane-adjacent and possibly newest PG layers (Meisner et al., 2013; Domínguez-Cuevas et al., 2013). Perhaps CwlO activity is more heavily regulated because it cleaves nascent PG to ensure its proper incorporation in the PG mesh, while the role of LytE is confined to expanding outer layers of PG in wild-type cells. Future experiments that crosslink LytE or CwlO to the PG strands they digest should allow detection of which strands are cleaved by each enzyme.

How does loss of *tkmA* bypass the requirement for *lytE* in a *ΔtagTV* mutant background (Fig 7)? TkmA encodes a Wzz (also called PCP) domain protein (Paysan-Lafosse et al., 2023). Wzz proteins form a membrane-anchored octameric tube that protrudes into the extracellular space between the cell membrane and PG, through which nascent glycopolymers are synthesized (Tocilj et al., 2008; Morona et al., 1995; Whitfield et al., 2020). In the biofilm-competent *B. subtilis* strain 3610, *tkmA* is required for biofilm formation in conjunction with its homolog *epsA* (Kiley and Stanley-Wall, 2010; Gerwig et al., 2014). In this strain background, the *ΔtkmA* biofilm defect can be rescued by overexpression of *epsA* (Gao et al., 2015). However, loss of *epsA* had no obvious effect in *ΔtagTUV* mutants, suggesting either that EPS is not the primary polymer processed by TkmA or that *tkmA* has an alternate function that impacts CwlO activity. Besides its presumed glycan-processing function, TkmA also promotes auto-phosphorylation of the DNA-binding protein PtkA, which then phosphorylates (and presumably, regulates the activity of) a number of substrates that may produce alternate PG-anchored polymers (Petranovic et al., 2007; Mijakovic et al., 2003; Mijakovic et al., 2004; Mijakovic et al., 2005; Derouiche et al., 2016; Jers et al., 2010). At this time, however, it is not possible to separate the envelope polymer processing function of TkmA from its indirect role in activating other polymer-synthesizing enzymes. In the future, labeling *ΔtkmA* cells to identify changes in polymer type or density might uncover changes that explain the primary function of *tkmA*.

It remains unclear how the polymers proposed to be synthesized by TmkA inhibit CwlO. Perhaps these hypothetical polymers physically interact with FtsEX or SweDC, affecting conformational changes that would allow CwlO to become active. Or the polymers may physically interfere with the CwlO enzymatic domain. Less direct models are also possible: for example, membrane-anchored polymers could create changes in the membrane that impact cytoskeletal organization or transmembrane PG synthesis machinery. It is also unclear whether these polymers play a role in maintaining periplasmic osmolarity, where they might aid in folding or activation of important extracellular protein complexes, including those that build PG. Further experiments are required to discover the precise nature and function of the inhibitory polymers.

Why are *ΔtagTUV* mutants viable? Cells mutated for *tagO*, which completely lack the earliest step of the enzymatic pathway required to make WTAs, are viable, though slow growing. In contrast, cells lacking later steps are completely inviable, due to trapping C55-P-linked intermediates that deplete the carrier lipid pool (D’Elia et al., 2006). LCP enzymes are essential in some Gram-positive organisms due to sequestering of C55-P (Malet-Villemagne et al., 2023; Wu et al, 2014). However, in *B. subtilis*, *ΔtagTUV* cells were able to grow, albeit at a very slow rate. I therefore propose that the depletion of the free C55-P pool because some of the polymers are hydrolyzed. This model suggests that there may be C55-P or a feedback loop that prevents polymer synthesis when polymers are not efficiently transferred. In *S. aureus*, WTAs are released from the membrane in the absence of all three LCP enzymes, suggesting that an enzyme that can cleave accumulated polymers might exist (Chan et al., 2013).

Under stressful conditions, bacteria must make a dramatic decision: lyse and die, or pause growth and wait for better conditions. If lysis is chosen, PG hydrolases – autolysins – degrade the PG layer and release the cytoplasm, killing the cell. If pausing is chosen, however, PG hydrolases must be inhibited until PG synthesis resumes. This work suggests that some polymers, when accumulated at the membrane, instead of being transferred, can inhibit a specific hydrolase. Inhibiting hydrolases might preserve the PG layer, maintaining the integrity of the cell until better conditions arrive.

## Materials and Methods

### General methods and strain construction

All *B. subtilis* strains were derived from the prototrophic strain PY79 (Zeigler et al., 2008). Cells were grown in LB medium at 37°C and then diluted to a mid-log phase titer (OD_600_ = 0.5) for all experiments. *B. subtilis* strains were constructed using plasmids, intact genomic DNA, or PCR-amplified fragments of genomic DNA introduced via a 1-step competence method (Dubnau and Davidoff-Abelson, 1971). Antibiotic concentrations were used at: 100 μg/ml spectinomycin, 5 μg/ml chloramphenicol, 10 μg/ml tetracycline, 10 μg/ml kanamycin and 1 μg/ml erythromycin. A list of strains, plasmids, and oligonucleotide primers used in this study can be found in Table S1, Table S2, and Table S3 in the supplemental material. The details of plasmid construction are included in Table S2.

### Fluorescence microscopy

Exponentially growing cells were harvested and concentrated by centrifugation at 4000 × g for 1.5 min, resuspended in 1/10 volume LB medium, and then immobilized on 2% (wt/vol) agarose pads containing LB medium. Fluorescence microscopy was performed on a Nikon Ti inverted microscope equipped with a Plan Apo 100×/1.4 Oil Ph3 DM phase contrast objective, an Andor Zyla 4.2 Plus sCMOS camera, and Lumencore SpectraX LED Illumination. Images were acquired using Nikon Elements 4.3 acquisition software. Venus was imaged using a Chroma ET filter cube for yellow fluorescent protein (49003) with an exposure time of 800 ms; mCherry was visualized using a Chroma ET filter cube (49008) with an exposure time of 100 ms; Image processing was performed using FIJI software (version 2.14.0/1.54f) (Schindelin et al., 2012).

### Immunoblot analysis

Immunoblot analysis was performed as described previously (34). Briefly, for each culture, 1 ml of cells, normalized to OD_600_ = 0.5, was collected and the cell pellet resuspended in 50 μl lysis buffer (20 mM Tris [pH 7.0], 10 mM MgCl2, 1 mM EDTA, 1 mg/ml lysozyme, 10 μg/ml DNase I, 100 μg/ml RNase A, 1× protease inhibitor cocktail (Roche)). The cells were incubated at 37°C for 10 min followed by addition of an equal volume sample buffer (0.25 M Tris [pH 6.8], 4% SDS, 20% glycerol, 10 mM EDTA) containing 10% β-mercaptoethanol. Samples were heated for 15 min at 65°C prior to loading. Proteins were separated by SDS-PAGE on 12.5% polyacrylamide gels, electroblotted onto nitrocellulose membranes (Thermo) and blocked in 3% bovine serum albumin (BSA) in phosphate-buffered saline (PBS) with 0.5% Tween 20. The blocked membranes were probed with anti-His (1:4,000) (Genscript) or anti-ScpB (1:10,000) (34) diluted into 3% BSA in 1× PBS with 0.05% Tween 20. The anti-His antibody was detected using anti-mouse IgG (Bio-Rad); anti-ScpB was detected using anti-rabbit IgG (Bio-Rad), and the Clarity Western ECL Blotting Substrate chemiluminescence reagent as described by the manufacturer (Bio-Rad). Signal was detected using an Azure 400 Imager (Azure Biosystems).

### Spot dilutions

Late-log cultures were normalized to OD_600_ = 1.5 and 10-fold serial dilutions were generated. 5 μL of each dilution were spotted onto LB-agar with or without supplemented IPTG. Plates were incubated at 37 °C overnight and photographed the next day.

## Acknowledgements

I would like to thank David Rudner for initiating this project and for generously offering his time and laboratory resources. Members of the Bernhardt-Rudner super-group past and present supplied much needed advice. Paula Montero Llopis and the HMS Microscopy Resources on the North Quad (MicRoN) core offered invaluable advice on microscopy. Fred Ausubel provided critical reading of the manuscript.

Support for this work comes from NIH grant 5R35GM145299 awarded to David Z. Rudner and 5F32GM146400 to Jennifer D. Cohen.

---

**Figure S1.**
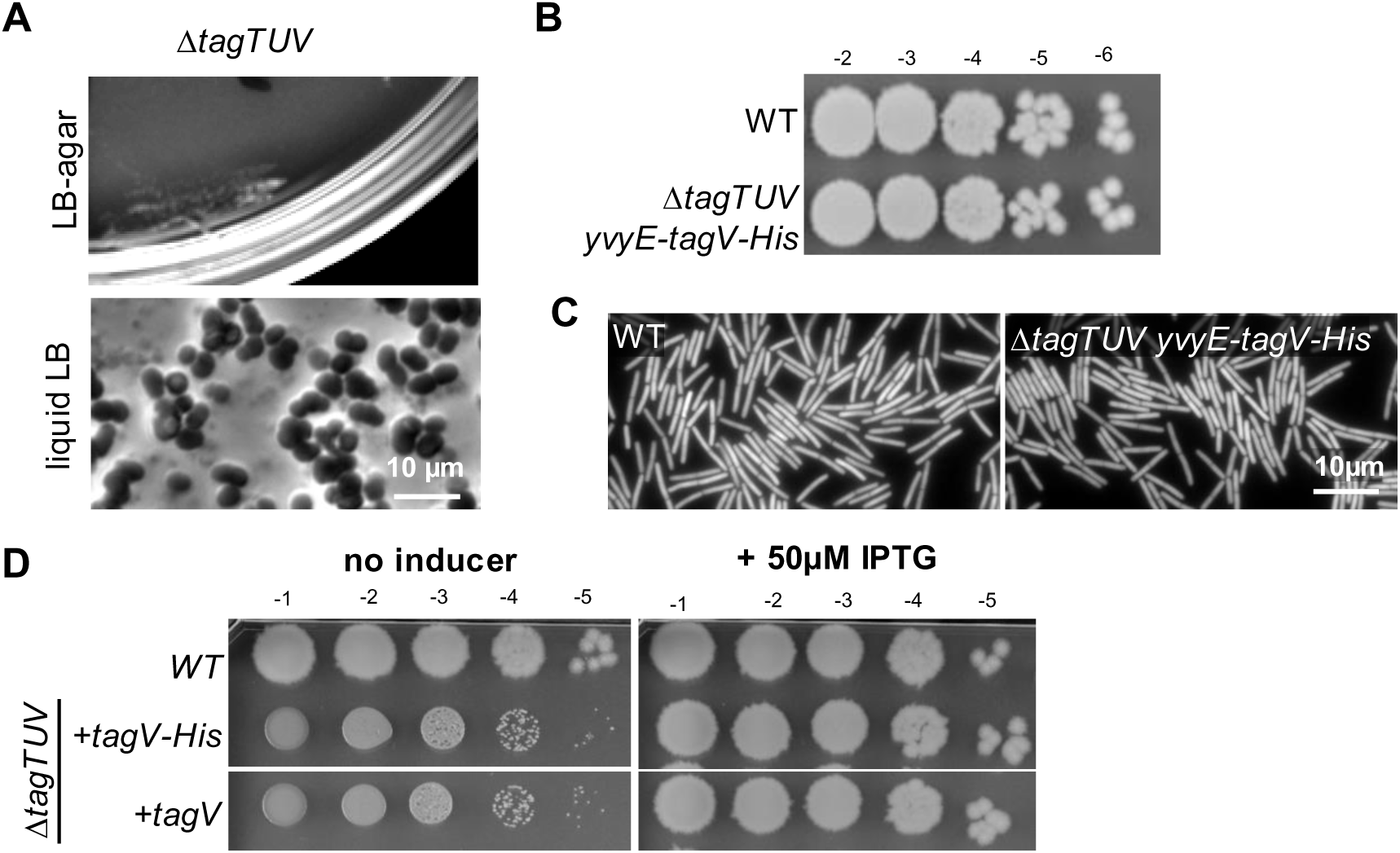
LCP mutants are viable, and TagV is sufficient for normal growth. A) Cells lacking all three LCP enzymes grew very slowly on LB-agar and had a rounded morphology when imaged under brightfield microscopy. B) Spot dilutions of the indicated strains. *yvyE-tagV-His* is expressed under its endogenous promoter from a non-native genomic locus (*yvbJ*) in cells deleted for *tagT* and *tagU*. C) Images of cells in mid-log growth expressing *sacA::Pveg-mCherry* to label cytoplasm. Rod-like cell morphology is visible in both WT cells and cells expressing *yvyE-tagV-His* as the sole LCP gene. D) Spot dilutions of the indicated strains in the presence and absence of IPTG. Representative plates from 3 independent replicates are shown.

**Figure S2.**
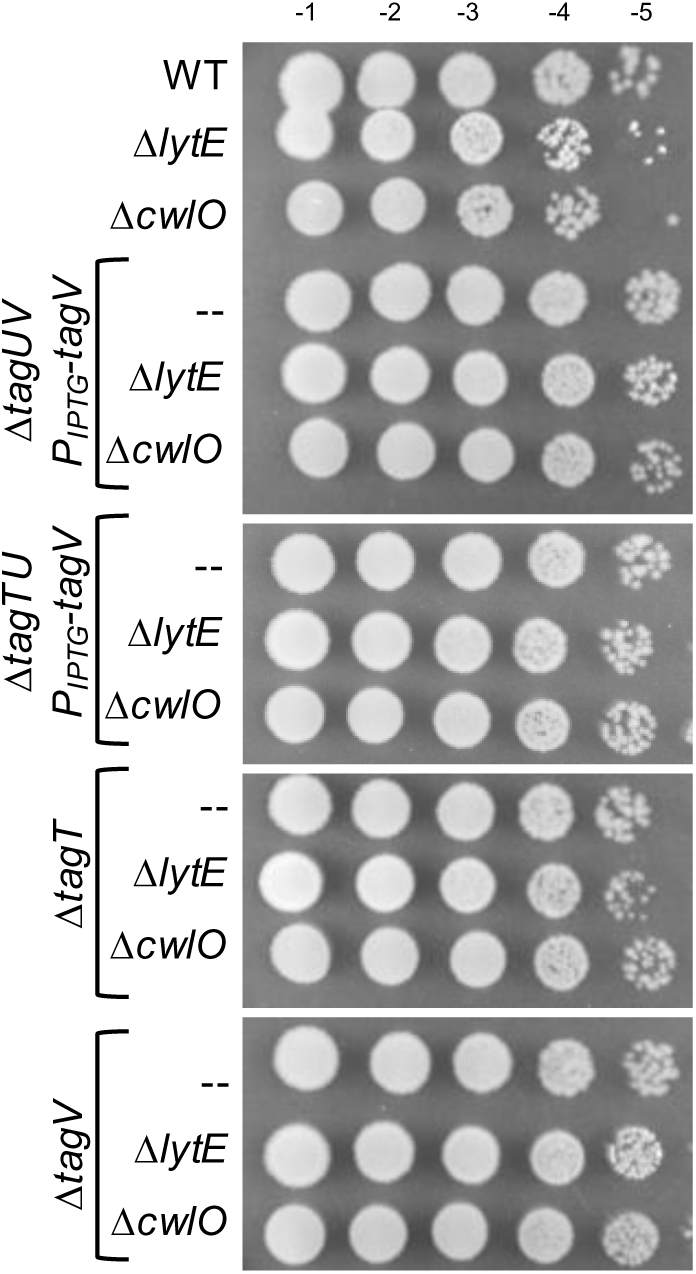
*lytE* is not synthetic sick with either *tagT* or *tagV* alone. Spot dilutions of the indicated strains on LB-agar. All strains grew at levels similar to WT.

**Figure S3.**
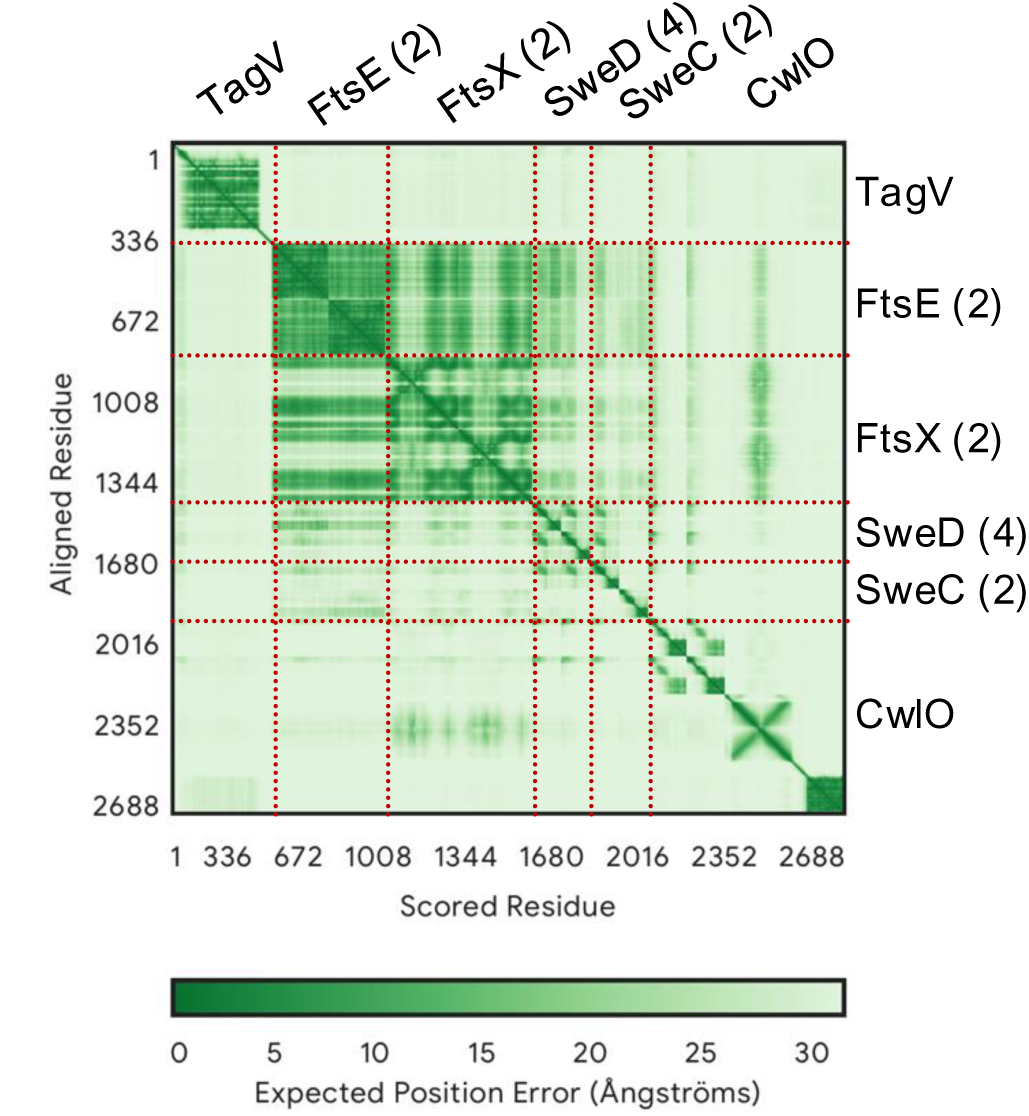
TagV is not predicted to interact with FtsEX-SweDC-CwlO. Alphafold3 predicts high error (low confidence) in interactions between TagV and any known CwlO complex proteins. In contrast, some interaction is predicted between FtsX, FtsE, SweD, and CwlO. Numbers indicate the number of proteins modeled; for example, 2 molecules of FtsE are modeled in this graph, after stoichiometry determined in Brunet et al., 2019.

**Figure S4.**
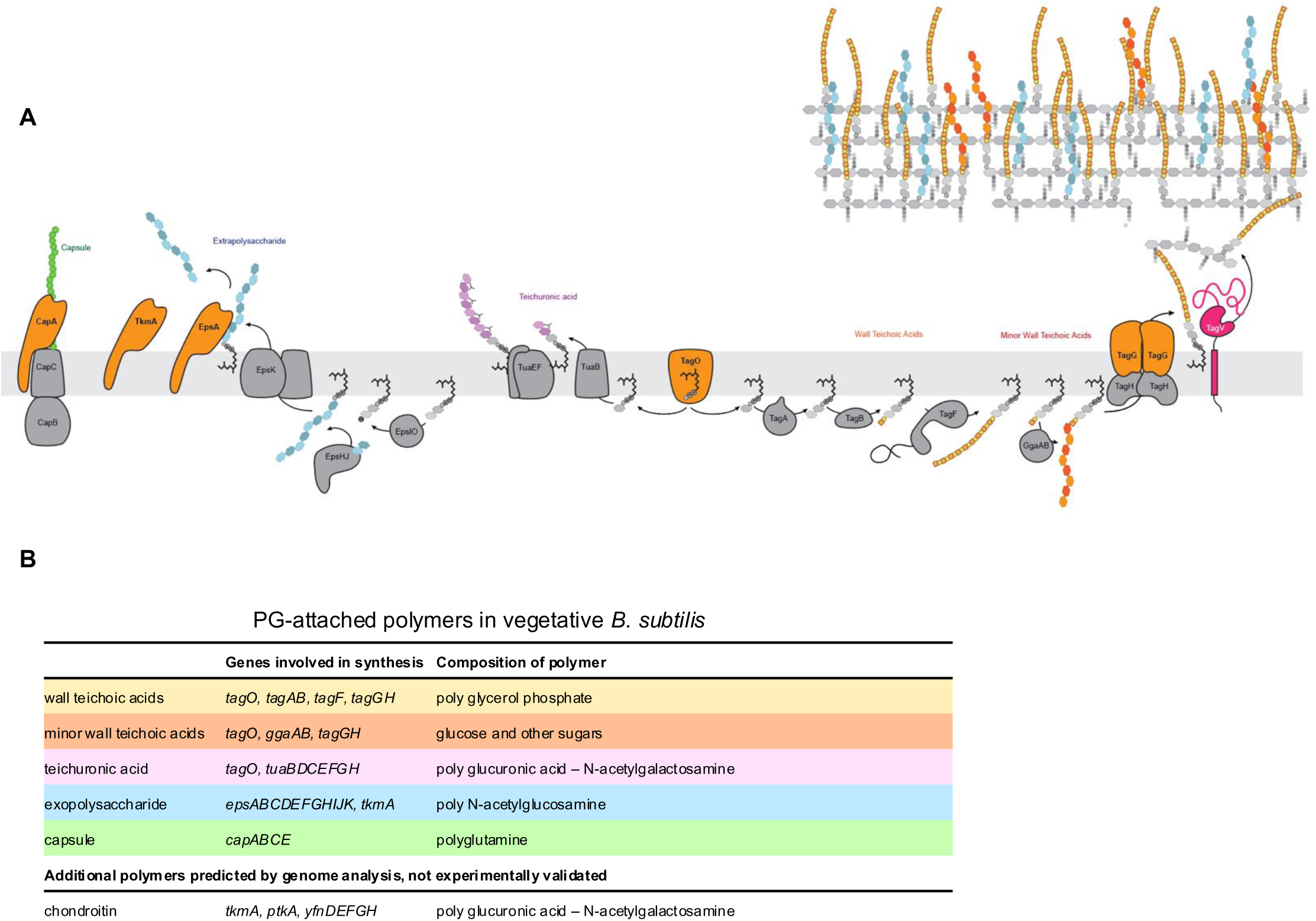
Pathways for synthesis of PG-anchored or secreted polymers in *B. subtilis*. A) Diagram of proposed biosynthetic pathways for PG-attached polymers. Polymers are colored as follows: green, capsule; blue, exopolysaccharide; purple, teichuronic acid; orange, minor teichoic acids; yellow, wall teichoic acids. Proteins removed via genetic deletion in this work are colored orange. PG is represented as a series of gray polymers above the membrane on the right. B) A list of all polymers colored to reflect the diagram above, with their main biosynthetic enzymes and compositions. After Baddiley, 1970; Soldo, 2002; Marvasi et al., 2010; Morikawa et al., 2006; Freymond et al., 2006.

## Notes

### Competing Interest Statement

The authors have declared no competing interest.

### Summary of Updates

The title was edited from "Glycopolymer transferases promote peptidoglycan hydrolysis in Bacillus subtilis" to "Evidence that glycopolymer transferases promote peptidoglycan hydrolysis in Bacillus subtilis".

